# Haplotracker: a web application for simple and accurate mitochondrial haplogrouping using short DNA fragments

**DOI:** 10.1101/2020.04.23.057646

**Authors:** Kijeong Kim, Yoonyeong Kim, Dong-han Kim, Chulhwan Kwon, Kyung-yong Kim

## Abstract

Mitochondrial DNA (mtDNA) haplogrouping is widely used in population genetics, forensics, and medical research to study evolutionary questions in human populations, examine degraded remains, and search for mtDNA-associated diseases. Next-generation sequencing methods have become a revolutionary tool for mtDNA analysis, particularly for degraded DNA, but they remain costly, laborious, and time-consuming. If an accurate and simple haplogroup-tracking tool were available, haplogrouping could be performed easily and rapidly. Here, we present Haplotracker, a web application for highly accurate and simple haplogroup tracking using a few sequence fragments. Haplotracker offers a unique user-friendly interface and estimates highly probable haplogroups from control-region sequences using a novel algorithm based on Phylotree Build 17 and our haplotype database (n=118,869). Haplotracker provides a simple, novel HG tracking solution, which was established through repeated blind simulation tests. It narrows down potential haplogroups and identifies their differential coding-region variants to confirm the haplogroups or to track sub-haplogroups. Haplotracker gives detailed information on sample variants, including their frequency in a large mtGenome database, which may give researchers an insight into common, rare, and potentially pathogenic mutations. It also offers a conserved region mapping tool for PCR primer design for successful tracking. Haplotracker produced high haplogroup prediction accuracy using 8,216 control-region Phylotree-provided sequences. It estimated top-ranking haplogroups with a higher concordance rate (56.6%, *p*<0.0001) than the similar tools MitoTool (29.4%) and HaploGrep 2 (33.9%). The significance persisted up to Rank 30. Haplotracker accurately estimated super-HGs from 94% of the control-region sequences at Rank 1. Further evaluation of the accuracy with 46,322 control-region sequences was significant. Laboratory application of Haplotracker to an ancient DNA extract demonstrated its practical usefulness. These results highlight the potential for the use of our web application as an alternative to full genome sequencing for easy haplogrouping, which may be useful in related fields. Free access: https://haplotracker.cau.ac.kr.

**Author summary:** Mitochondrial haplogroup (HG) classification is required in the search for answers to evolutionary questions about human populations, in the investigation of forensic samples, and in the search for disease-associated mutations. The sequencing of mitochondrial DNA (mtDNA) at the genome level is now frequently used due to the falling cost of next-generation sequencing. It offers accurate and detailed analysis of mtDNA variation, but it is still costly, laborious, and time-consuming. If an accurate and simple HG tracking tool were available, haplogrouping could be performed easily and rapidly. Here, we developed Haplotracker, a simple and accurate HG tracking web application with a novel algorithm and a novel tracking tool. The highly accurate prediction performance of Haplotracker was demonstrated in a series of tests. Using only control-region sequences, it accurately predicted HGs in more than half of the total Phylotree-provided mtGenome sequences and approximately 80% up to Rank 5; for super-HGs, it accurately predicted 94% at Rank 1 and more than 98% up to Rank 3. We demonstrated simple tracking for HG confirmation using Haplotracker. These results show that mtDNA haplogrouping with our web application may be useful in related fields.

## Introduction

Mitochondrial DNA (mtDNA) haplogroup (HG) assignment, known as mtDNA haplogrouping, is widely used in population genetics, molecular anthropology, and forensics [1–14]. It is used to study evolutionary questions in human populations, examine degraded remains, and search for disease-associated mutations [15]. Haplogrouping also helps in the comparison and communication of genetic variation [16] and is important for sequence data quality control (QC) [13]. Haplogrouping is usually conducted using computer software that automatically assigns the HG of a given mtDNA sequence based on Phylotree, the tree of global mtDNA variation, because manual haplogrouping is time-consuming and error-prone [17]. The most recent version of Phylotree is Build 17, which is available at https://www.phylotree.org [18].

Haplogrouping of fresh DNA samples, from which the sequence information for large mtDNA fragments can be obtained, is much easier than that of degraded DNA samples, such as those obtained from forensic specimens or human remains. Their DNAs are present in a very small amount and are shortly fragmented, often containing exogenous materials that may inhibit or interfere with PCR [19, 20]. These characteristics render haplogrouping infeasible. As a result, the many sequence fragments often need to be sequenced. Next-generation sequencing (NGS) methods with enrichment technology have thus become a revolutionary tool for the study of mtDNA sequence variation, particularly from degraded DNA samples at the mtDNA genome (mtGenome) level [21–23], because conventional Sanger sequencing is labor-intensive, expensive, and time-consuming for complete mtGenome sequencing [24]. For the purpose of haplogrouping, however, HG prediction using several control-region (CR) sequence fragments and subsequent confirmation using coding-region information may be a rapid and simple alternative, reducing the costs, labor, and time required. The CR sequences of mtDNA, which comprise three hypervariable (HV) regions – HV region 1 (HV1) (positions 16024–16365), HV region 2 (HV2) (positions 73–340), and HV region 3 (HV3) (positions 438–574) – are common targets used for the haplogrouping of degraded samples because their polymorphic properties enable them to be used to discriminate haplotypes or HGs [25–29]. However, coding-region information is often required for refined haplogrouping when CR sequences are used [13].

Currently available computer-assisted tools for mtDNA haplogrouping include mtDNAmanager [29], MitoTool [30], HaploGrep 2 [31], EMPOP [32, 33], HAPLOFIND [34], and James Lick’s mtHap (https://dna.jameslick.com/mthap/). In particular, HaploGrep 2, MitoTool, and EMPOP have become widely used [5–12] because they represent powerful servers with complete mtGenome sequences and provide useful additional functions for mtDNA variation analysis. However, while they can estimate the most probable HGs from CR sequences, they do not suggest additional solutions to confirm the results or determine sub-HGs. These solutions are important because HG estimates using CR sequences require further confirmation given that they are less accurate than the use of mtGenome sequences and often result in multiple HG estimates. For example, the determination of the HG of an mtDNA sequence (GenBank accession number: AY289059), which is defined as HG “O” by Phylotree, depends on whether the mtGenome or CR sequence is used. When the mtGenome sequence is used, it is consistently assigned to “O” by MitoTool, HaploGrep 2, EMPOP, and Haplotracker. In contrast, when the CR sequence is used, the most probable HG is identified as “N9b2a” by MitoTool and “N9b2” by the other servers. MitoTool did not predict “O” as an option at all, while the other tools ranked it second.

These results demonstrate that subsequent confirmation is required for HG prediction when using CR sequences. This confirmation and sub-haplogrouping are usually conducted using differential variants in the coding region. If many indeterminable HGs are highly ranked in the haplogrouping process, it will be laborious and complicated. A simple tracking method is thus required to definitively and accurately determine the HGs because higher prediction accuracy results in a less laborious process. Several papers have raised concerns about accurate HG prediction. Bandelt et al. remarked that most journal articles in forensic genetics presented haplogrouping in rudimentary or incorrect ways [13], while Röck et al. also pointed out the problem of automated solutions that did not provide reliable and unbiased HG predictions, especially with partial mtDNA sequences [32]. In contrast, MitoTool has been reported to be outstanding in terms of accuracy [30], and HaploGrep 2 has increased its accuracy by improving its underlying algorithm [31].

In the present study, we develop Haplotracker, a web application that offers simple and highly accurate tracking of mtDNA HGs using CR and coding-region sequence fragments. Haplotracker offers a unique user-friendly interface with input options for multiple sequence fragments without the need to input the ranges from an mtDNA sample. The server first uses several short sequence fragments in the CR of mtDNA to produce a list of the closest-ranked HGs using a novel algorithm based on Phylotree definitions and our haplotype database (DB). To confirm the predicted HGs, Haplotracker employs a novel HG tracking tool. It minimizes the number of tests for HG tracking by narrowing down the HGs through integration to their most recent common ancestor (MRCA) HGs. It then identifies highly specific and conserved differential variants among these HGs. To confirm the HGs or identify the sub-HGs, researchers are guided to re-track the haplogrouping with additional sequence fragments across the positions of the differential variants. The server also provides manual options and tools for researchers to select and compare HGs to narrow down and differentiate HGs. These include HG selection options, options to determine the level of sub-HGs ranging from MRCAs to terminal sub-HGs, access to the HG DB, and a separate tool for HG differentiation.

Ambiguous HGs, even those in the top ranks, can be compared to determine differential variants. To re-track the HGs, the presence of HG-differential variants in the sample DNA needs to be examined. This requires PCR amplification of DNA fragments located across the variant positions. Hybridizable primer designs for the fragments are essential for successful PCR, particularly for degraded DNA. Haplotracker provides a mapping tool showing the other variants present across the target variants to survey possible conserved regions for the primers to minimize the chance of potential primer mismatches. In addition, Haplotracker provides information about the sample variants, including their frequencies observed in a large mtGenome DB, which may be helpful for researching commonly found, HG-defining, rare, and/or potential disease-associated variants within a sample. Because the QC of mtDNA sequences is also important to ensure sequencing reliability [35, 36], Haplotracker includes QC tools for the detection of possible artificial recombinations [37] and phantom mutations [38] in a dataset.

To evaluate the performance of Haplotracker in HG prediction using CR sequences, we examined haplogrouping with CR sequences from 8,216 mtGenome sequence examples provided by Phylotree and 46,322 CR sequences from mtGenome sequences downloaded from GenBank. We then compared the haplogrouping results with those of MitoTool and HaploGrep 2.

## Results

### Input

We designed a user-friendly interface for the input of multiple sequence fragments or variant profiles from a sample (S1 Fig.). It includes DNA sequence or variant fields for the fragments and the following additional fields: sample name, an option for the score DB (Haplotracker or HelixMTdb), super-HG level, rank group level, an option for accepting “N” nucleobases as variants, and selection buttons for the control or coding region in which the dominant part of the fragment sequence is positioned. Researchers can input up to 100 sequence fragments regardless of any overlap in the separate fields without needing to input the ranges of the fragments. Researchers can increase the number of input fields for the fragments by clicking the “+” buttons beneath the field or reduce them by clicking the “-“ buttons. The rank group level (1–4) is used to set the lowest HG rank group to display. “N” nucleobases in sequences are interpreted as “not sequenced” by default but can be selected to be interpreted as “A, C, G, or T” by ticking the checkbox. In the latter case, more than four tandem-repeated “N”s are interpreted as “not sequenced.” Ticking the button “Control” or “Coding” for the fragment region is important for correct processing. All of the sequence fragments in a sample are aligned with the revised Cambridge Reference Sequence (rCRS) using the implemented Gotoh algorithm. There is no need to input the nucleotide positional ranges of the fragments.

Once the variants and ranges of the fragments are obtained using the first tool, a second tool, “HG tracking by fragment variant profiles,” can be used with these data rather than the sequences themselves for HG tracking (S2 Fig.).

### HG tracking flowchart

A simple HG tracking flowchart for Haplotracker was constructed based on repeated blind simulation tests with many Phylotree-defined samples with different ranks ranging from 1 to 50 and samples ranked over 50 belonging to every major HG. First, Haplotracker predicts HGs with CR sequences, particularly using the three HV region sequences. Re-tracking of the HGs is required to find diagnostic differential variants in the coding regions for HG confirmation or sub-haplogrouping. For the simple tracking of HGs, we suggest the strategy outlined in Fig. 1. If more than one HG is present in Rank Group 1, we propose narrowing down the HGs in this group first. This is based on our observation that most of the definitive HGs (95%) that were identified by Haplotracker from the CR regions (n=8,216) of Phylotree mtGenome sequences were found in Rank Group 1 (Table 1). If more than one HG is scored (i.e., has a score >0), it is preferable to first track the HGs with scores. This is also based on our results showing that most of the definitive HGs (99.6%) in Rank Group 1 were found among the scored HGs within this group. They can be differentiated using specific variants offered by the server. Further narrowing down of the HGs can be conducted if needed. If there are too many HGs to track, differentiation between the top five scored HGs or rank group MRCAs can be conducted as an alternative approach. Tracking is accompanied by the acquisition of the fragment sequences across the variant positions using PCR amplification. Tracking can be repeated several times to verify the HGs or track sub-HGs. Tracking is completed following the final confirmation of the MRCA of Rank Group 1 by differentiating between the MRCAs of the rank groups.

**Fig. 1.**
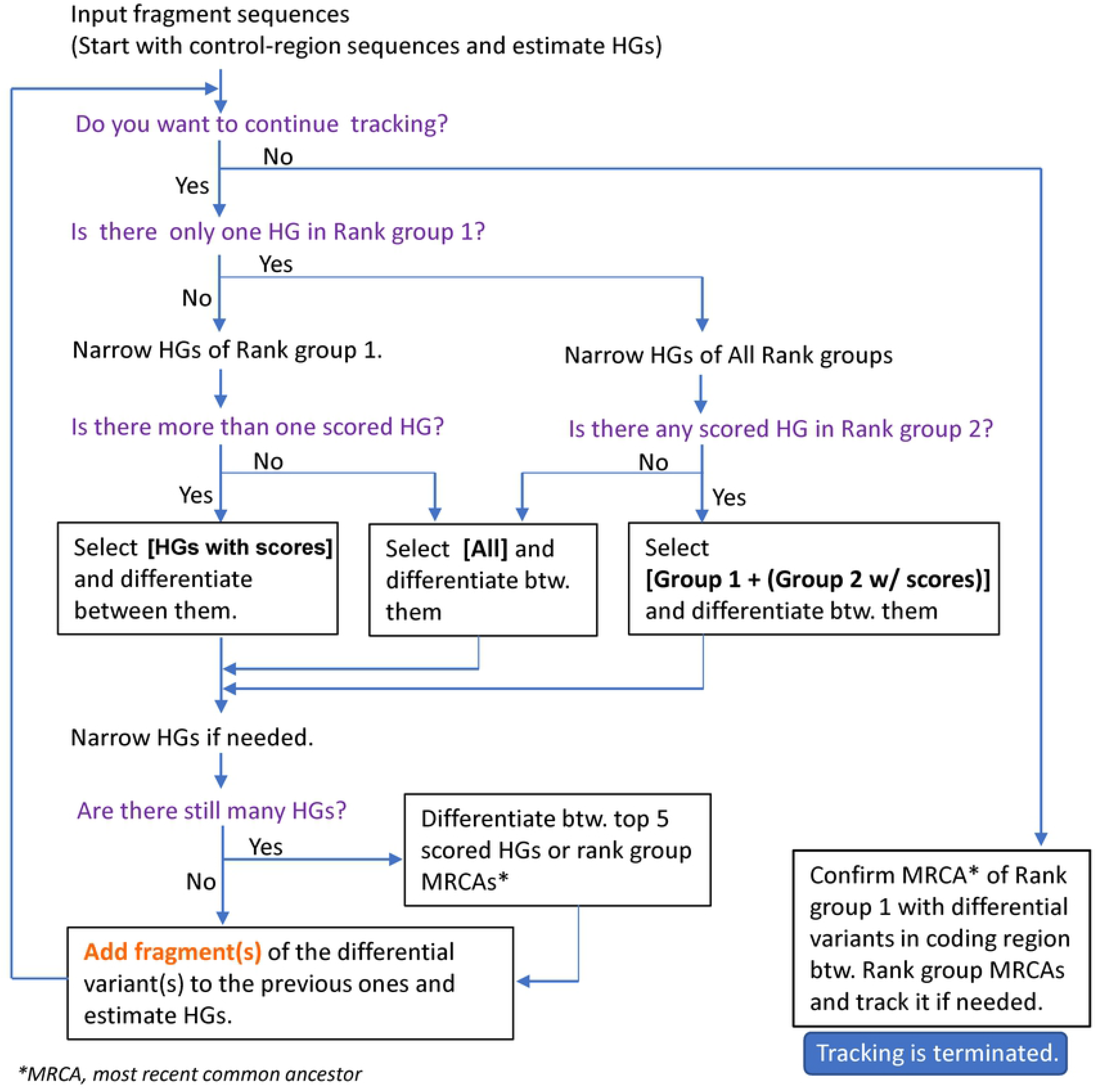
HG tracking flowchart describing a strategy for the use of Haplotracker to track HG confirmation or sub-haplogrouping. If there is more than one HG in Rank Group 1 after HG estimation using CR sequences, the narrowing down of the HGs in Rank Group 1 is attempted first. If there is more than one scored HG (score >0), it is preferable to track the HGs with scores first. They can be differentiated using specific variants on the server. Further narrowing down of the HGs can be conducted if needed. If there are too many HGs to track, differentiation between the top five scored HGs or rank group MRCAs can be conducted. Tracking can be repeated several times to confirm the HGs or track sub-HGs. Tracking ends by verifying the MRCA of Rank Group 1 by differentiating between the rank group MRCAs. HG, haplogroup; CR, control-region; MRCA, most recent common ancestor.

**Table 1.** HG prediction accuracy of Haplotracker using CR sequences of mtGenome sequences from Phylotree.

| Phylotree-defined HG | Rank Group 1 | Rank Group 2 | Total (n=8,216) |
| --- | --- | --- | --- |
| Identified | 7,804 (95.0%) | 392 (4.8%) | 8,196 (99.8%) |
| Score of 0 only in the rank group | 4,143 (50.4%) | 2 (0.0%) | 4,145 (50.5%) |
| Score >0 only in the rank group | 1,488 (18.1%) | 2 (0.0%) | 1,490 (18.1%) |
| Score of 0, Score >0 in the rank group | 2,173 (26.4%) | 388 (4.7%) | 2,561 (31.2%) |
| Score >0 from the row above | 2,164 (99.6%) | 342 (88.1%) | 2,506 (97.9%) |
| Not identified |  |  | 20 (0.2%) |

### Output

#### Track by sequence

This first main tool displays the sample variant information extracted from the inputted sequence fragments and the list of highly probable HG estimates (Table 2, Fig. 2). When researchers choose HelixMTdb as the score DB, Haplotracker displays the HG estimates at super-HG Level 1 (L3, M, G, D, N, R, HV, H, T, U, etc.) because the DB provides the variant frequencies of super-HGs. When researchers restrict the super-HG level in Haplotracker, the server displays the HG estimates accordingly. Without the restriction, it displays all of the probable estimates of the HGs present in Phylotree. Haplotracker then provides options for HG confirmation or sub-HG tracking along with details in a new window with the sample name on the browser tab. The results can be downloaded as an HTML file.

**Table 2.**
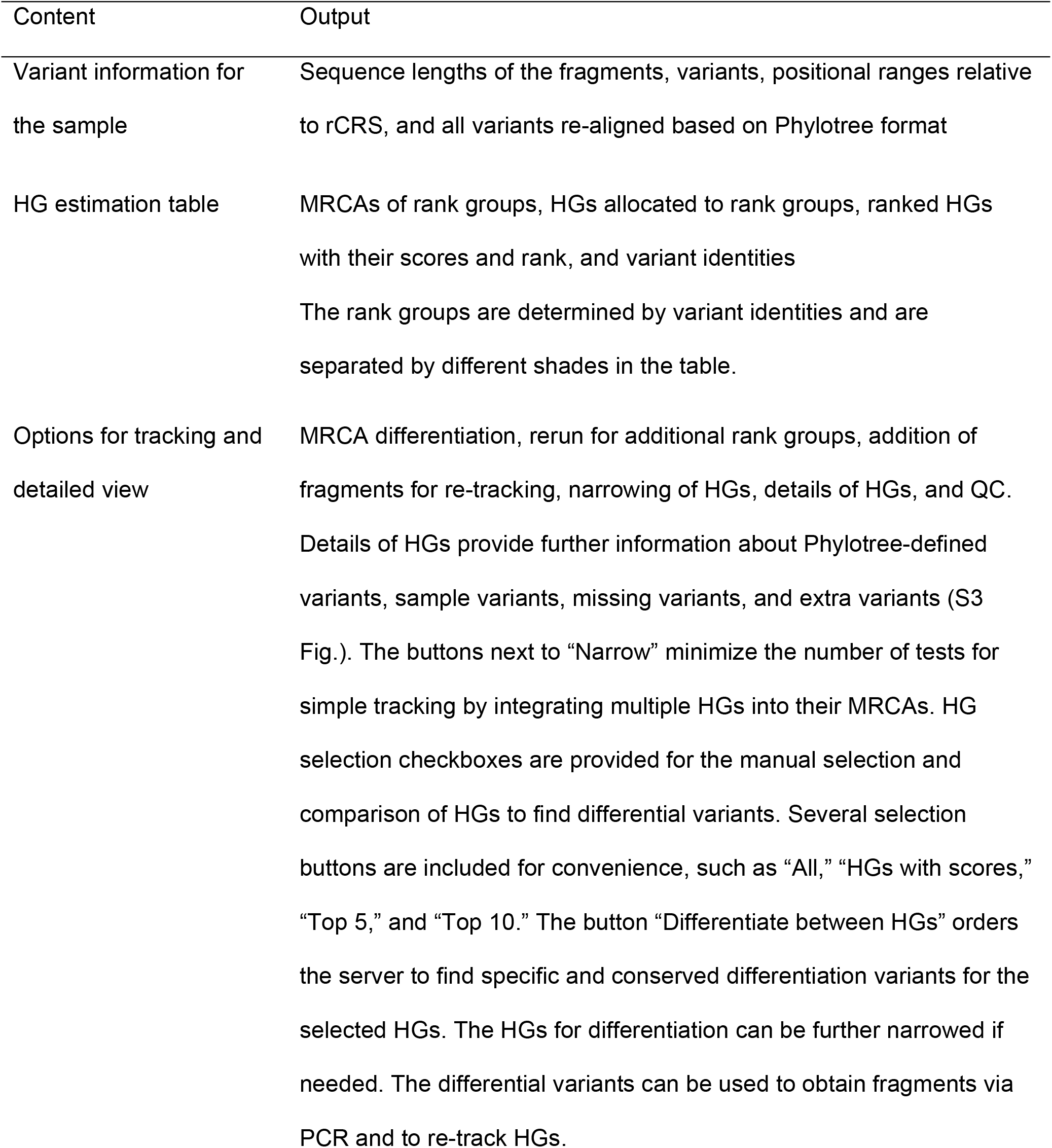
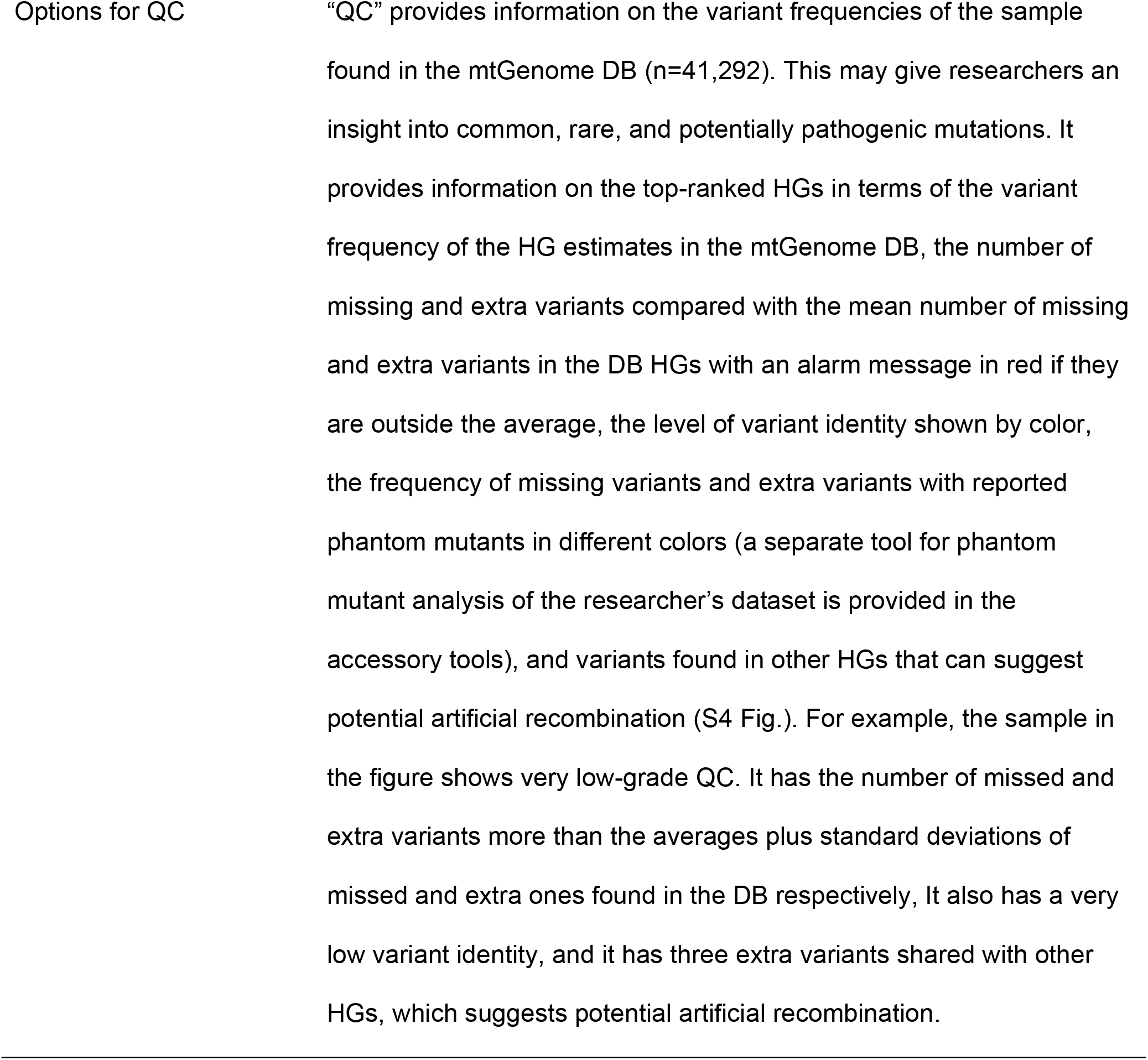
HG tracking output from Haplotracker.

**Fig. 2.**
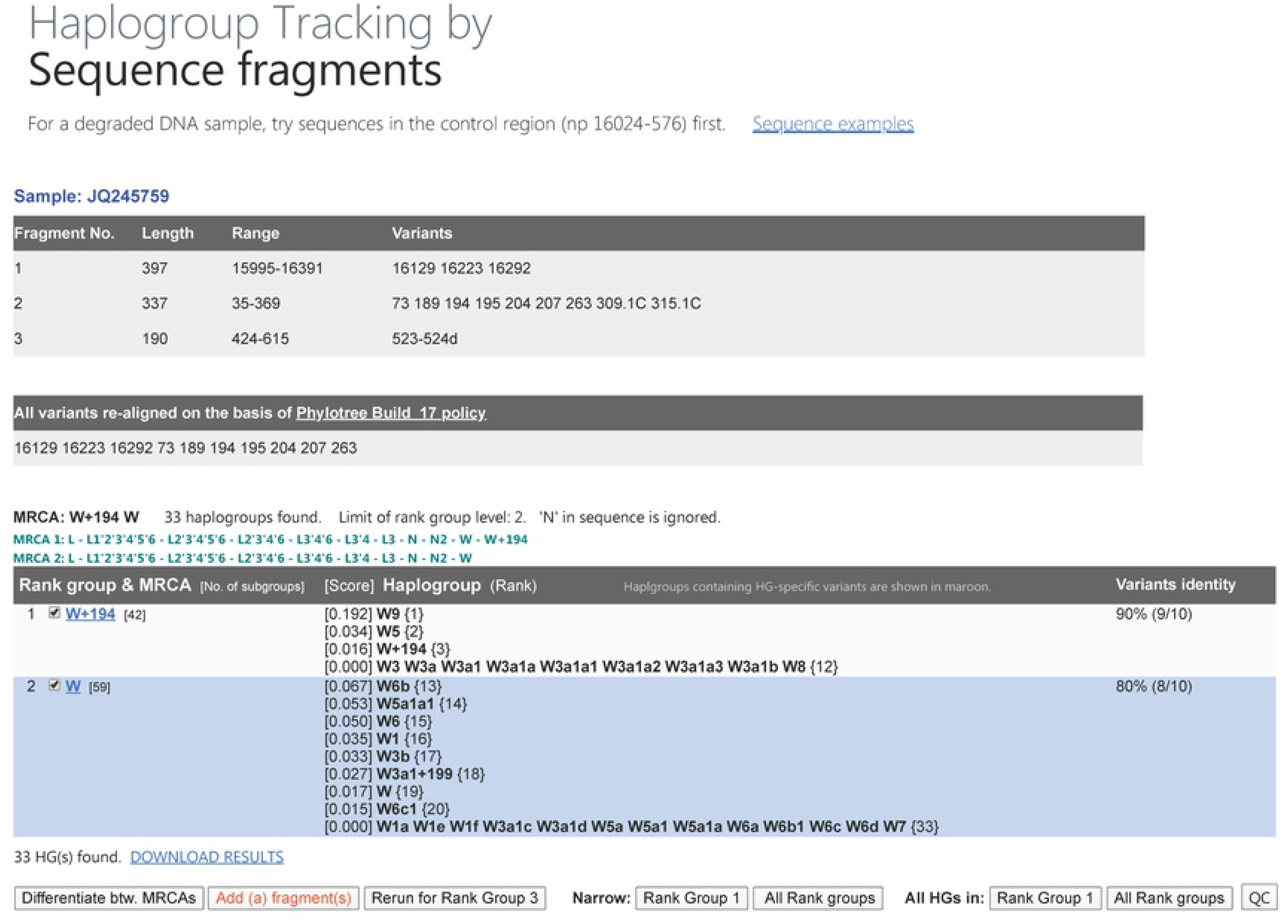
HG estimation results showing variant information for the sample, rank groups, MRCAs, and ranks of the estimated HGs. The tracked HGs (boldface) were first divided into rank groups with their MRCA (first column), which was determined by their variant identity (third column). The HGs were listed (second column) in order of ranking (shown in parentheses to the right of the HG), as determined by their scores (shown in brackets to the left of the HG). In this example, there are 12 tracked HGs in Rank Group 1 and 21 in Rank Group 2. HG W9 was the most highly ranked with a score of 0.192 and was predicted to be the highest for the given sample sequences. Options for simple tracking for HG confirmation or sub-haplogrouping are provided along with options for details and QC information. HG, haplogroup; MRCA, most recent common ancestor.

#### Track by variant

This tool provides the same functions as the first tool but with the input of the fragment variant profiles produced by the tool above. It allows for the use of variant information obtained from assays, such as SNaPshot or probe-based real-time PCR assays, in addition to sequencing. It can also be used for simulation tests prior to the input of experimental data into Haplotracker.

#### Conserved region map

This tool searches the conserved regions in the given range and HGs, displaying them and variant positions in a variant map, with the HGs at the positions in a table form (S5 Fig.). When a diagnostic variant for the confirmation of a predicted HG is selected, PCR of the fragment across the position of the variant is required to confirm the presence of the variation. Well-designed primers are essential for successful PCR. The primers should be perfectly hybridized with the template DNA of the sample, especially for degraded DNA that is only available in small quantities. This tool helps researchers find conserved regions for primer hybridization with a given range and set of HGs. Even if there are no conserved regions found under the given conditions and no other options, researchers can design primers with degenerated sequences relying on information from the variants suggested by this tool. Under the given range with or without HG restriction, this tool finds all variants present within the range and presents them graphically and the HGs for every variant.

#### Accessory tools

The “HG Database” tool helps researchers to explore the HG variants defined by Phylotree. An HG of interest is entered in the HG field and the lowest sub-HG level to be listed is specified. The server then returns all of the HGs from the HG entered to the lowest level of its subgroups, with their HG-defining variants, differentiation variants between the listed HGs, highly specific variants, and non-conserved variants. The “HG Differentiation” tool compares HGs entered by the researchers. It has an interface and functions similar to those of “HG Database” but multiple independent HGs can be inputted. These tools were designed to help researchers find highly specific diagnostic variants mostly in the coding region to further confirm the predicted HG or to determine the specific HG among a group of predicted HGs. The highest specific variants of an HG (maroon) are those found only in the HG and its subgroups, if any; highly specific variants (teal) are those found in the HG but also found in a small number of other HGs; non-conserved variants (gray) are those not found in the HG subgroups; and differential variants (boldface) are those that can be used to specifically determine the HG among the listed HGs. The highly specific variants are listed in a separate column of the output table with the number of cross-presenting HGs in brackets (a smaller number means it is more specific). The non-conserved variants in the separate column are listed with the number of non-conserved subgroups in brackets (a larger number means it is less conserved). The recommended variants for further HG testing will be the highest specific variants as the first priority, highly specific variants as the second priority, and highly conserved variants as the third priority. The “Phantom Mutants” tool searches for these in a dataset among the extra variants that are not used for HG definition by Phylotree. The output results are the same as those for HaploGrep 2. The “Haplogrouping with Complete mtGenome Sequences (MtGenome Haplogrouping)” tool assigns the sample the HG based on a complete mtGenome sequence. This is not the primary goal of this server. We use this tool for Haplotracker’s mtGenome-based HG assignments and compare them with those of HaploGrep 2. The “Variant Format Conversion” tool allows researchers to produce variant profiles in different formats, including Phylotree, HaploGrep 2, EMPOP, and MitoTool. It should be noted that the variant profiles can be server-specific, so they are not fully interchangeable for haplogrouping. The variants obtained by a certain web server can only be used for haplogrouping by the same server. “Major HG-specific Variants” can help researchers to locate a major HG in the tree of Phylotree Build 17.

### Comparison of web servers

We compared the haplogrouping performance of MitoTool, HaploGrep 2, and Haplotracker using different sources of mtDNA sequences. The results of the top-ranked HGs are summarized in Table 3, and detailed results of other ranks can be found in the supporting data.

**Table 3.**
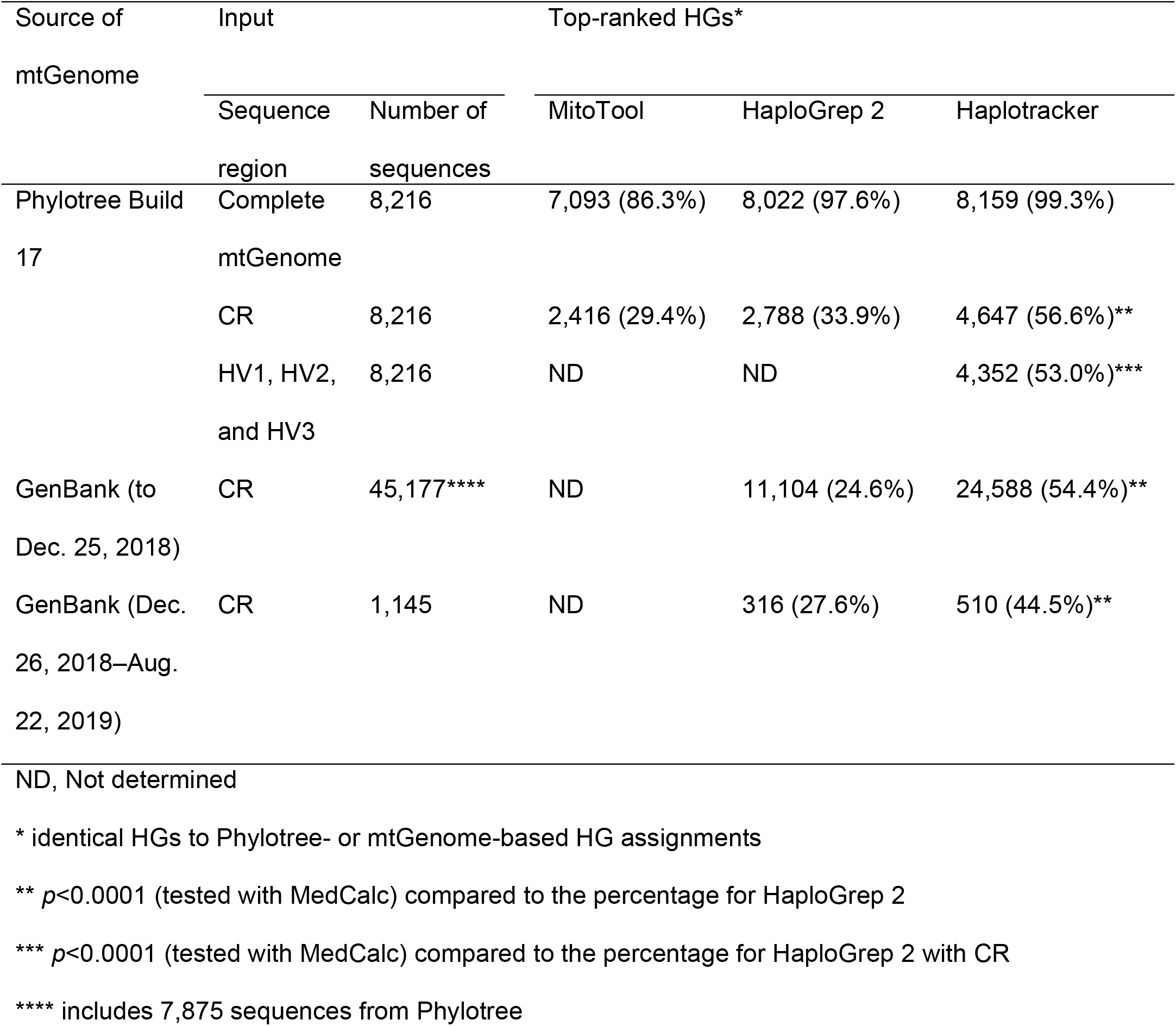
Comparison of haplogrouping performance among servers.

#### HG estimation with Phylotree mtGenome sequences (n=8,216)

Haplotracker predicted HGs with the highest concordance rate (99.3%) with Phylotree HGs, with concordance lower for MitoTool (86.3%) and HaploGrep 2 (97.6%) (S1 Table). There was no significant difference in HG assignment between HaploGrep 2 and Haplotracker, but significant (*p*<0.0001) differences were found between MitoTool and HaploGrep2 and between MitoTool and Haplotracker.

#### HG estimation with Phylotree CR sequences (n=8,216)

Haplotracker predicted HGs with high accuracy (Table 4, Fig. 3, S2 and S3 Tables, and S6 Fig.). It predicted the top-ranking HGs with a higher concordance rate (56.6%, *p*<0.0001) with Phylotree HGs than did MitoTool (29.4%) and HaploGrep 2 (33.9%). The significantly higher accuracy of Haplotracker was observed from Ranks 1 to 30. With Haplotracker, approximately 80% of the CR sequences were assigned to concordant HGs in Ranks 1–6. The rate of unidentified Phylotree HGs in any rank was lower with Haplotracker (0.2%, *p*<0.0001) than with MitoTool (56.1%) and HaploGrep 2 (6.7%).

**Table 4.**
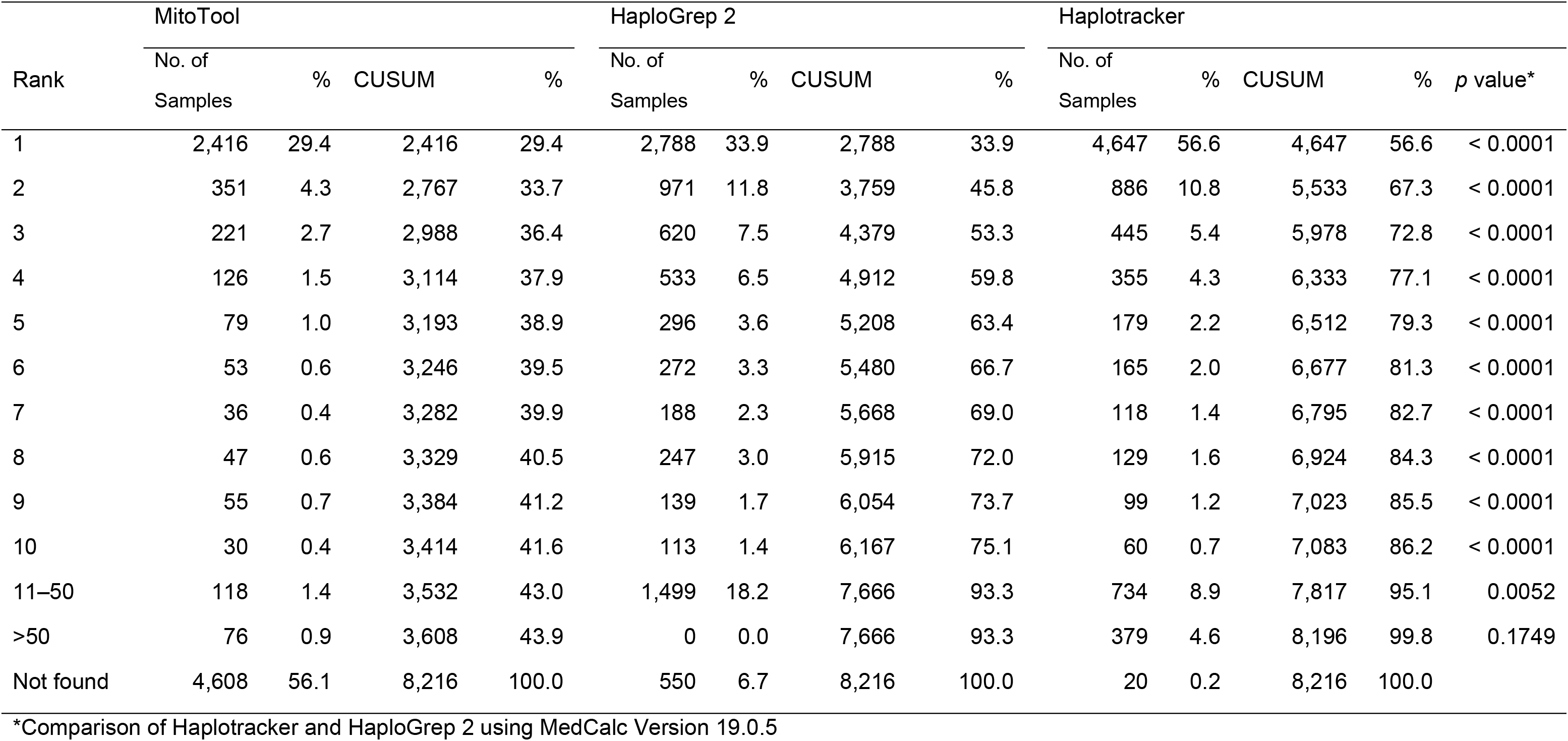
Comparison of HG prediction concordance rates for MitoTool, HaploGrep 2, and Haplotracker with Phylotree-defined HGs using only the CR of mtGenome sequences (n=8,216) provided by Phylotree mtDNA tree Build 17.

| Rank | MitoTool |  |  |  | HaploGrep 2 |  |  |  | Haplotracker |  |  |  |  |
| --- | --- | --- | --- | --- | --- | --- | --- | --- | --- | --- | --- | --- | --- |
|  | No. of Samples | % | CUSUM | % | No. of Samples | % | CUSUM | % | No. of Samples | % | CUSUM | % | p value* |
| 1 | 2,416 | 29.4 | 2,416 | 29.4 | 2,788 | 33.9 | 2,788 | 33.9 | 4,647 | 56.6 | 4,647 | 56.6 | < 0.0001 |
| 2 | 351 | 4.3 | 2,767 | 33.7 | 971 | 11.8 | 3,759 | 45.8 | 886 | 10.8 | 5,533 | 67.3 | < 0.0001 |
| 3 | 221 | 2.7 | 2,988 | 36.4 | 620 | 7.5 | 4,379 | 53.3 | 445 | 5.4 | 5,978 | 72.8 | < 0.0001 |
| 4 | 126 | 1.5 | 3,114 | 37.9 | 533 | 6.5 | 4,912 | 59.8 | 355 | 4.3 | 6,333 | 77.1 | < 0.0001 |
| 5 | 79 | 1.0 | 3,193 | 38.9 | 296 | 3.6 | 5,208 | 63.4 | 179 | 2.2 | 6,512 | 79.3 | < 0.0001 |
| 6 | 53 | 0.6 | 3,246 | 39.5 | 272 | 3.3 | 5,480 | 66.7 | 165 | 2.0 | 6,677 | 81.3 | < 0.0001 |
| 7 | 36 | 0.4 | 3,282 | 39.9 | 188 | 2.3 | 5,668 | 69.0 | 118 | 1.4 | 6,795 | 82.7 | < 0.0001 |
| 8 | 47 | 0.6 | 3,329 | 40.5 | 247 | 3.0 | 5,915 | 72.0 | 129 | 1.6 | 6,924 | 84.3 | < 0.0001 |
| 9 | 55 | 0.7 | 3,384 | 41.2 | 139 | 1.7 | 6,054 | 73.7 | 99 | 1.2 | 7,023 | 85.5 | < 0.0001 |
| 10 | 30 | 0.4 | 3,414 | 41.6 | 113 | 1.4 | 6,167 | 75.1 | 60 | 0.7 | 7,083 | 86.2 | < 0.0001 |
| 11–50 | 118 | 1.4 | 3,532 | 43.0 | 1,499 | 18.2 | 7,666 | 93.3 | 734 | 8.9 | 7,817 | 95.1 | 0.0052 |
| >50 | 76 | 0.9 | 3,608 | 43.9 | 0 | 0.0 | 7,666 | 93.3 | 379 | 4.6 | 8,196 | 99.8 | 0.1749 |
| Not found | 4,608 | 56.1 | 8,216 | 100.0 | 550 | 6.7 | 8,216 | 100.0 | 20 | 0.2 | 8,216 | 100.0 |  |

\*Comparison of Haplotracker and HaploGrep 2 using MedCalc Version 19.0.5

**Fig. 3.**
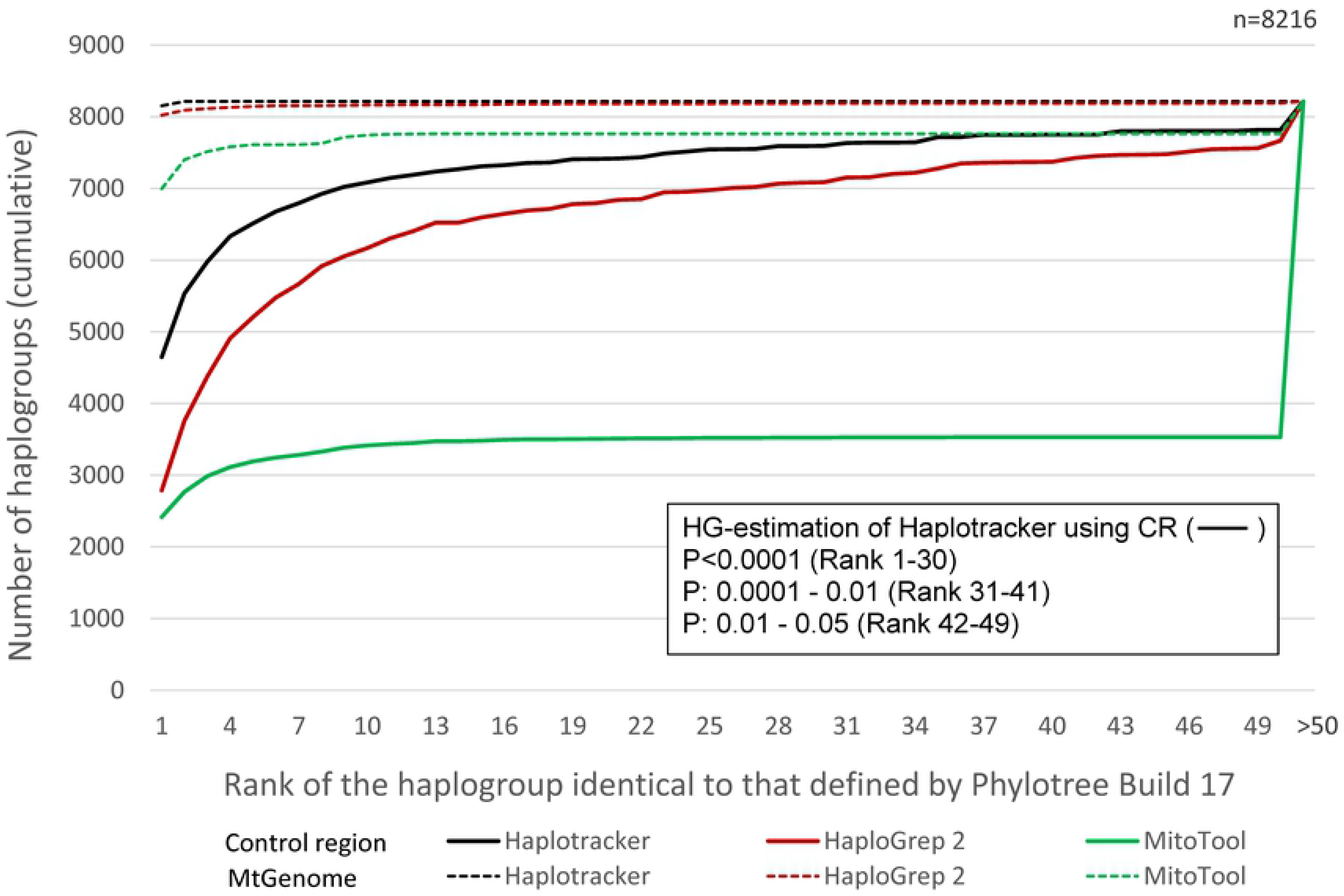
Comparison of HG assignment with the complete mtGenome and with mtDNA CR sequences using MitoTool, HaploGrep 2, and Haplotracker. These data are based on the cumulative number of estimated HGs identical to Phylotree HGs at a rank as tested with the mtGenome and CR of 8,216 mtGenome sequences provided by Phylotree. HG, haplogroup; mtGenome, mitochondrial genome; mtDNA, mitochondrial DNA; CR, control-region.

#### Further extensive evaluation (n=45,177, CR)

Haplotracker estimated the top-ranking HGs concordant with mtGenome-based HGs at a rate higher than that of HaploGrep 2 (54.4% and 24.6%, respectively; *p*<0.0001) (S4 and S5 Tables) using CR sequences. Haplotracker also showed significantly higher accuracy for Ranks 1 to 49. The rate of unidentified HGs using Haplotracker (1.0%, *p*<0.0001) was lower than that of HaploGrep 2 (6.1%). HaploGrep 2 experienced a server error with 43 sequences.

#### Evaluation with new sequences (n=1,145, CR)

Haplotracker estimated the top-ranking HGs concordant with mtGenome-based HGs at a rate higher than that of HaploGrep 2 (44.5% and 27.6%; *p*<0.0001) (S6 Table) using CR sequences. Haplotracker also exhibited significantly higher accuracy for Ranks 1 to 9. Haplotracker’s rate of unidentified HGs (1.0%, *p*=0.0114) was lower than that of HaploGrep 2 (5.0%).

#### Haplotracker HG estimation with HV region sequences

Using HV region sequences, Haplotracker estimated top-ranking HGs at a high concordance rate (53.0%) with Phylotree HGs. Its accuracy was lower (*p*=0.0019) than prediction using CR sequences (56.6%) but higher (*p*<0.0001) than the accuracy of HaploGrep 2 using CR sequences (33.9%). The accuracy of Haplotracker with HV sequences was lower (*p*<0.05) than the estimation with CR sequences for Ranks 1–4 but higher (*p*<0.001) than that of HaploGrep 2 in Ranks 1–30 (S2 and S7 Tables).

### Super-HG estimation of Haplotracker using CR sequences

Using the CR from the Phylotree mtGenome sequences, Haplotracker exhibited high prediction rates for the super-HGs. More than 93% of the total sequences (n=8,216) were concordantly estimated for Rank 1 and 99.8% up to Rank 3 (Table 5). Most of the super-HGs had a prediction rate of over 95% for Rank 1, while the H, HV, and V haplogroups had a relatively low prediction rate (<78%). However, including Ranks 2 and 3, these haplogroups had a prediction rate of more than 98%, with most of the rest having a 100% prediction rate.

**Table 5.** Super-HG estimation by Haplotracker using CR sequences from Phylotree Build 17.

| HG |  | Rank 1 |  | Rank 2 |  |  | Rank 3 |  |  |
| --- | --- | --- | --- | --- | --- | --- | --- | --- | --- |
| Name | No. of | No. of | % | No. of | CUSUM | % | No. of | CUSUM | % |
|  | Samples | Samples |  | Samples |  |  | Samples |  |  |
| L0 | 201 | 201 | 100 |  | 201 | 100 |  | 201 | 100 |
| L1 | 123 | 123 | 100 |  | 123 | 100 |  | 123 | 100 |
| L5 | 15 | 15 | 100 |  | 15 | 100 |  | 15 | 100 |
| L2 | 188 | 188 | 100 |  | 188 | 100 |  | 188 | 100 |
| L6 | 4 | 4 | 100 |  | 4 | 100 |  | 4 | 100 |
| L4 | 21 | 21 | 100 |  | 21 | 100 |  | 21 | 100 |
| L3 | 277 | 276 | 99.6 | 1 | 277 | 100 |  | 277 | 100 |
| M | 923 | 903 | 97.8 | 17 | 920 | 99.7 | 3 | 923 | 100 |
| C | 231 | 231 | 100 |  | 231 | 100 |  | 231 | 100 |
| Z | 36 | 36 | 100 |  | 36 | 100 |  | 36 | 100 |
| E | 55 | 55 | 100 |  | 55 | 100 |  | 55 | 100 |
| G | 106 | 99 | 93.4 | 3 | 102 | 96.2 | 4 | 106 | 100 |
| Q | 52 | 52 | 100 |  | 52 | 100 |  | 52 | 100 |
| D | 501 | 473 | 94.4 | 7 | 480 | 95.8 | 21 | 501 | 100 |
| N | 171 | 171 | 100 |  | 171 | 100 |  | 171 | 100 |
| I | 104 | 104 | 100 |  | 104 | 100 |  | 104 | 100 |
| W | 100 | 100 | 100 |  | 100 | 100 |  | 100 | 100 |
| Y | 24 | 24 | 100 |  | 24 | 100 |  | 24 | 100 |
| A | 260 | 260 | 100 |  | 260 | 100 |  | 260 | 100 |
| O | 3 | 2 | 66.7 | 1 | 3 | 100 |  | 3 | 100 |
| S | 9 | 7 | 77.8 | 2 | 9 | 100 |  | 9 | 100 |
| X | 128 | 128 | 100 |  | 128 | 100 |  | 128 | 100 |
| R | 231 | 223 | 96.5 | 5 | 228 | 98.7 | 1 | 229 | 99.1 |
| HV | 155 | 119 | 76.8 | 26 | 145 | 93.5 | 8 | 153 | 98.7 |
| V | 105 | 69 | 65.7 | 36 | 105 | 100 |  | 105 | 100 |
| H | 1,607 | 1,252 | 77.9 | 82 | 1,334 | 83.0 | 258 | 1,592 | 99.1 |
| J | 395 | 395 | 100 |  | 395 | 100 |  | 395 | 100 |
| T | 385 | 385 | 100 |  | 385 | 100 |  | 385 | 100 |
| F | 146 | 145 | 99.3 | 1 | 146 | 100 |  | 146 | 100 |
| B | 510 | 510 | 100 |  | 510 | 100 |  | 510 | 100 |
| P | 32 | 30 | 93.8 | 1 | 31 | 96.9 |  | 31 | 96.9 |
| U | 801 | 799 | 99.8 | 2 | 801 | 100 |  | 801 | 100 |
| K | 317 | 316 | 99.7 | 1 | 317 | 100 |  | 317 | 100 |
| Total | 8,216 | 7,716 | 93.9 |  | 7,901 | 95.7 |  | 8,196 | 99.8 |

### Laboratory application of Haplotracker

The laboratory-obtained sequence data from an ancient DNA sample (MNX3) were used to evaluate the performance of Haplotracker in experimental research. The sequence electropherograms of the amplified products appeared clean, as shown in S8 Table. First, with three inputted sequence fragments of the CR (HV1 and HV2), Haplotracker predicted the HG of MNX3 to be “U2e1a1” as the top rank (Table 6). The MRCA of Rank Group 1 was predicted to be “U2e1,” and this group had nine sub-HGs. Of these, HG U2e1a1 was top-ranked with the highest score. To confirm U2e1a1, we followed the steps suggested in the haplogrouping flowchart (Fig. 1). The first question in the flowchart is “Do you want to continue tracking?” The answer was “Yes.” Following the arrow, the second question is “Is there only one HG in Rank Group 1?” The answer was “No” because there were nine HGs in Rank Group 1. We followed the subsequent instruction and narrowed down the HGs of Rank Group 1 by clicking the button [Rank Group 1]. As a result, the HGs were narrowed down from nine to six. The next arrow arrives at the question “Is there more than one scored HG?” The answer was “Yes” because there were three scored HGs. We followed the arrow and arrived at “Select [HGs with scores] and differentiate between them.” The results showed that there were three HGs (U2e1a1, U2e1a, and U2e1) and one HG differential variant “3116” for U2e1a1. We followed the next arrow and arrived at “[Add fragment(s)] of the differential variant(s) to the previous ones and predict HGs.” We added the sequence of the fragment for the “3116” coding-region position that was obtained from the PCR experiment in the present study and pressed the submit button. The result screen showed that, in the upper table, the sample MNX3 had the variant “3116,” which is the coding-region variant of U2e1a1. There was no extra variant in the coding-region fragment. The lower table showed that U2e1a1 was top-ranked and U2e1a1c was ranked second. This means that the most probable HG predicted using only CR sequences was confirmed by the coding-region variant. The MRCA of Rank Group 1 was also U2e1a1, which is U2e1 with the CR sequences above. We continued tracking by following the chart. We clicked the buttons to narrow and differentiate between the two HGs. Haplotracker displayed the differential variant “10127 (U2e1a1c).” We added the PCR-obtained sequence fragment for 10207 to the previous inputs and pressed submit. The results showed that MNX3 did not have the 10207 variant and that only U2e1a1 was in Rank Group 1. Haplotracker thus successfully discriminated U2e1a1 from U2e1a1c. We additionally confirmed this HG with PCR-obtained sequences for 11197 (U2e1a) and 6045 (U2e) proposed by Haplotracker. The results showed that the HG prediction was consistently accurate.

**Table 6.** Laboratory results for the application of Haplotracker to ancient DNA haplogrouping.

| Track No. | Sequence fragments* | Rank Group 1 |  | Score | Rank | Variants found |
| --- | --- | --- | --- | --- | --- | --- |
|  |  | MRCA | HGs |  |  |  |
| 1 | 15972–16193 (HV1),<br>16182–16410 (HV1),<br>15–389 (HV2) | U2e1 | U2e1a1 | 0.101 | 1 | 16051 16093 16129C<br>16189 16362 73 152<br>217 263 340 |
|  |  |  | U2e1a1c | 0.015 | 2 | Same as above |
|  |  |  | U2e1a | 0.013 | 3 | Same as above |
|  |  |  | U2e1 | 0.000 | 4 | Same as above |
|  |  |  | U2e1a1a | 0.000 | 9 | Same as above |
|  |  |  | U2e1c |  |  |  |
|  |  |  | U2e1c1 |  |  |  |
|  |  |  | U2e1d U2e1g |  |  |  |
| 2 | + 3050-3197 (**3116**<br>for U2e1a1) | U2e1a1 | U2e1a1 | 0.101 | 1 | 16051 16093 16129C<br>16189 16362 73 152<br>217 263 340 **3116** |
|  |  |  | U2e1a1c | 0.015 | 2 | Same as above |
| 3 | + 10010-10179<br>(**10207** for U2e1a1c) | U2e1a1 | U2e1a1 | 0.101 | 1 | 16051 16093 16129C<br>16189 16362 73 152<br>217 263 340 **3116** |
| 4 | + 11137-11281<br>(**11197** for U2e1a) | U2e1a1 | U2e1a1 | 0.101 | 1 | 16051 16093 16129C<br>16189 16362 73 152<br>217 263 340 **3116**<br>**11197** |
| 5 | + 5978-6128<br>(**6045** for U2e) | U2e1a1 | U2e1a1 | 0.101 | 1 | 16051 16093 16129C<br>16189 16362 73 152<br>217 263 340 **3116**<br>**11197 6045** |

Boldface, differential variants in the coding region \*+, addition to sequence fragments of the previous track

In summary, we outlined an example of Haplotracker being employed in a laboratory experiment involving a degraded sample to evaluate its haplogrouping performance. With Haplotracker, we were able to determine and confirm the refined HG status of MNX3 (U2e1a1), which had been previously identified as U2e1 by a Neighbor-joining tree method, U2e by mtDNAmanager and U by Haplogroup Prediction Tool, and manually selected coding-region variants for U2 and U2e in a previous report [4].

We also used other web servers to haplogroup the same sequence fragments from MNX3. However, we were not able to use MitoTool or HaploGrep 2 because these tools do not support the input of multiple fragment sequences from a sample. EMPOP accepts the input of multiple fragments, but the input interface is not user-friendly. The precise ranges of all the fragments are also requested as input. It does not allow even partially overlapping fragments like the two HV1 fragments in this study. All of the ranges had to be separated by a space and entered into a single field, and likewise, all of the sequences had to be entered into another field that was too small for all of the sequence fragments to be seen at a glance. This required us to be very careful to avoid errors when entering multiple fragments and causing mix-ups among the ranges and sequences. The haplogrouping results with EMPOP using the HV1 and HV2 sequence fragments were U2e1 and U2e1f with MRCA U2e1 in Rank Group 1. U2e1a1a and others were found in Rank Group 2. Neither U2e1a1 nor U2e1a was found among the HGs displayed by EMPOP. Nonetheless, when we added the sequence fragment for “3116” proposed by Haplotracker, EMPOP predicted HG U2e1a1 as the top rank with U2e1a1 as the MRCA, as did Haplotracker.

## Discussion

A number of mtDNA servers provide unique and useful functions for mtDNA sequence variation analysis and haplogrouping. MitoTool provides four unique functions: 1) presentation of the variant location, the interspecies conservation index, and changes in amino acid status; 2) identification of potentially pathogenic mutations; 3) statistical analysis for HG distribution frequency; and 4) batch downloading of analytical output and mitochondrion-related data of interest. EMPOP provides a suite of software to support the analysis and interpretation of mtDNA sequence variation: 1) an HG browser representing the most recent Phylotree HGs in a convenient searchable format displaying their geographic locations graphically, 2) a tool that performs plausibility checks on an rCRS-coded data table, and 3) a tool that draws quasi-median networks that are useful for examining the quality of an mtDNA dataset. HaploGrep 2 rapidly treats mtDNA sequences on a mass scale for haplogrouping with a graphical display and handles high-throughput data that is compatible with NGS. It offers unique additional support, such as 1) immediate QC based on a generic rule-based system to detect artificial recombinants, missing variants, and rare/phantom mutations; 2) real-time graphical reports for multiple sequence alignment format, VCF format, and extended HG QC reports; 3) publication-ready phylogenetic trees for all input samples; and 4) an algorithm that increases accuracy and speed.

Haplotracker has several special haplogrouping features that distinguish it from other similar web applications in that it is able to accurately track HGs even from short DNA fragments, such as degraded DNA. Most web servers exhibit similar haplogrouping performance when using complete mtGenome sequences [31]; however, Haplotracker estimates HG with high accuracy using fragmented CR or HV region sequences. The present study demonstrated this difference with a high number of sequences from different sources. With CR sequences, Haplotracker provided estimates that were closest to the Phylotree-defined and mtGenome-based HG assignments. This higher HG estimation accuracy is believed to be due to our novel algorithm, which first separates the rank groups, which fulfills the requirements for Phylotree HG definitions, and then separates HG estimates restrictively inside each rank group using the scores produced from the private variant frequencies observed in the large dataset of haplotypes built into the server. Indeed, the value of private mutations for reliable haplogrouping has been previously noted [32].

Accurately estimating HGs can reduce the number of subsequent times HG confirmation must be conducted. Haplotracker helps researchers to select HG-specific and conserved coding-region fragments for confirmation. Haplotracker was designed with a user-friendly input interface for multiple raw sequence fragments without the need to know the fragment locations on rCRS, which is convenient and appropriate for testing multiple fragmented samples. The other servers do not support this. Haplotracker also provides simple HG tracking options to minimize the number of subsequent tests with the estimated HGs, which are generally numerous at a high rank when only CR sequences are used. These functions were successfully demonstrated in the present study by applying Haplotracker to a laboratory experiment with an ancient DNA sample in this study. To the best of our knowledge, there have been no reported web applications specifically designed with HG tracking tools for the simple confirmation and refinement of CR-based HGs from multiple fragmented DNA samples. Instead of CR sequencing, multiplex PCR methods for the identification of HG-determinative variants in coding regions have been reportedly used for degraded DNA [39, 40]. These methods may represent alternative approaches for haplogrouping. However, they have limitations for sub-haplogrouping in that they only assign super-HGs and do not suggest sub-HGs, which makes further tracking difficult.

The use of full mtGenome sequencing has become more common, and there are a number of excellent servers available for accurate haplogrouping using complete mtGenome sequences, such as EMPOP and HaploGrep 2. EMPOP exhibited higher accuracy with full mtGenome sequences than HaploGrep 2 and MitoTool (data not shown), while HaploGrep 2 delivered rapid, high-throughput results even with full mtGenome sequences. However, complete mtGenome sequencing is not always necessary for the purpose of haplogrouping depending on the sample. In many samples, only a few tracking tests are required to identify the HG as determined by mtGenome. In this study, Haplotracker was able to predict HGs identical to the mtGenome-based HGs for more than half of the full sequences provided by Phylotree using only the HV region sequences. Using only CR sequences, Haplotracker haplogrouped approximately 80% of Phylotree-provided mtDNA sequences up to Rank 5. A few more confirmation tests would allow mtGenome-based HGs to be identified. Even for low-ranked samples that may require many additional tests for HG differentiation, Haplotracker offers researchers an approach for the simple haplogrouping of these samples. Many of the very low-ranked samples are able to be haplogrouped using our HG tracking strategy (data not shown). Haplotracker was able to determine the Phylotree-defined super-HGs in more than 93% of the samples for Rank 1 and more than 98% up to Rank 3 using only CR sequences. This accurate prediction of super-HGs can also be utilized for a simple HG tracking strategy. Haplotracker outputs detailed information on the sample variant frequencies in a large mtGenome database in its system. This may give researchers an insight about common, rare, and/or potentially pathogenic mutations. An NGS method for CR sequencing and the use of targeted coding-region variants for global HGs and East Asian HGs have previously been adopted for the haplogrouping of degraded DNA [24]. Haplotracker may be applied to this method to simply estimate HGs using the data. Haplotracker can be a useful software application for the simple and accurate haplogrouping of degraded samples from various sources. Our server programs and hardware environment will be constantly updated in parallel with the future accumulation of mtDNA haplotype data.

### Methods

Haplotracker was coded using the active server page (ASP) script of Internet Information Services (IIS) Version 8.0 for Windows Server 2012 and Javascript. It was implemented using Gotoh, an open-source C implementation of the Gotoh algorithm, also known as the Needleman-Wunsch algorithm, with affine gap penalties (https://github.com/oboes/gotoh) to search the variants and nucleotide positions of the sample sequences relative to the rCRS [41, 42]. The variants were realigned according to the Phylotree format. The server works with most of the current versions of web browsers.

Haplotracker consists of three main tools and six accessory tools. The main tools include HG tracking by fragment sequences (Track by Sequence), HG tracking by fragment variant profiles (Track by Variant), and conserved region search for primer design (Conserved Region Map). HG tracking is the representative tool of our server. It extracts variants and ranges from the sample sequence fragments compared with the rCRS using the implemented Gotoh tool, and it tracks the most probable HGs based on the algorithm that we developed in this study.

### Algorithm

The HG of a sample was first tracked by comparing the sample variants with variant profiles of Phylotree-defined HGs (Phylotree HGs) (S1 Dataset). The highly probable HGs from the sample were listed in ordered rank groups. A rank group was determined by variant identity, the number of Phylotree-defined variants (Phylotree variants) of an HG present in the sample variants (Vp) minus the number of Phylotree variants of the HG missing (not present) from the sample variants (Vm) in all the ranges of the fragments, which was divided by the number of sample variants (Vs). The variant identities were calculated using all of the Phylotree HGs (n=5,434) using the equation below. The HGs with the highest variant identity were grouped as Rank Group 1, the second-highest as Rank Group 2, the third-highest as Rank Group 3, and the fourth-highest as Rank Group 4.

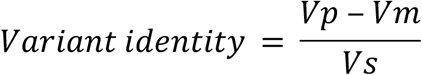

The example summarized in Table 7 and S7 Fig. consists of a sample with the variants “73 103 204 263 16092 16140 16189 16243 16278” in the sequence range of 16024–576 (CR). The HGs predicted in Rank Group 1 (B5b1, B5b, and B5b1a) had a variant identity of 77.8% (7/9), Vp = 7 (73 103 204 263 16140 16189 16243), Vm = 0 (no missing variants), and Vs = 9 (number of sample variants). Rank Group 2 had three HGs (B5b1c, B5b1c1, and B5b1c1a) with a variant identity of 66.7% (6/9), Vp = 7 (73 103 204 263 16140 16189 16243), Vm = 1 (152), and Vs = 9.

**Table 7.** Prediction of HGs with sample variants listed in order of rank group and rank.

| Rank Group | HG* | Phylo tree variant profile in CR | Sample variant profile | Missing variant(s) | Extra variant(s) | Variant identity | Score |
| --- | --- | --- | --- | --- | --- | --- | --- |
| 1 | B5b1 (1) | **73 103 204** | **73 103 204** |  | 16092 | 77.8% | 0.033 |
|  |  | **263 16140** | **263** 16092 |  | 16278 | (7/9) |  |
|  |  | **16189 16243** | **16140 16189** |  |  |  |  |
|  |  |  | **16243** 16278 |  |  |  |  |
|  | B5b (3) | **73 103 204** | **73 103 204** |  | 16092 | 77.8% | 0.000 |
|  | B5b1a (3) | **263 16140** | **263** 16092 |  | 16278 | (7/9) |  |
|  |  | **16189 16243** | **16140 16189** |  |  |  |  |
|  |  |  | **16243** 16278 |  |  |  |  |
| 2 | B5b1c (6) | **73 103 152** | **73 103 204** | 152 | 16092 | 66.7% | 0.000 |
|  | B5b1c1 (6) | **204 263** | **263** 16092 |  | 16278 | (6/9) |  |
|  | B5b1c1a (6) | **16140 16189** | **16140 16189** |  |  |  |  |
|  |  | **16243** | **16243** 16278 |  |  |  |  |

Boldface indicates those variants identical to the Phylotree variant profile, while underlining is used to indicate missing variants
\* Ranks are shown in parentheses.

The HGs were further ranked within each rank group by their scores. The scores were produced by the sum of the frequency rates of the HG carrying the extra and missing variants found in the haplotype DB (n=118,869) of Haplotracker (S2, S3, and S4 Dataset). The equation is as follows:

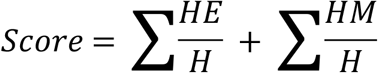

where HE is the frequency of the HG carrying the extra variant, HM is the frequency of the HG carrying the missing variant, and H is the frequency of the HG in the DB. The haplotype DB was constructed by repeating HG prediction runs three times with variant identity and regenerated scores using the mtDNA sequences downloaded from GenBank. Erratic sequences reported in previous papers were not used [13, 36, 38]. The DB contained 49,066 complete or partially complete genome sequences and 69,803 CR sequences.

For example, the top-ranked HG of the sample shown in Table 1 was predicted to be B5b1 with the highest score of 0.033. The sample had two extra variants, 16092 and 16278. The sum of the frequency rates of these extra variants of HG B5b1 is as follows:

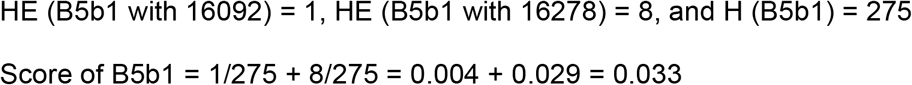

The ranks were determined in each rank group. This process differs from the algorithms employed by other servers. We first valued the variant identities more highly than the scores themselves. In addition to our DB, we integrated HelixMTdb (43) into our server to calculate scores based on the HGs and their private variant frequencies of HelixMTdb. This DB contains 15,035 unique variants from the mtGenomes of 196,554 unrelated individuals. However, it provides haplotype information for super-HGs only. Researchers can use HelixMTdb for HG estimation. Haplotracker uses the scores to rank HGs inside the rank group.

Narrowing down and differentiating the estimated HGs for further tracking were based on the tree structures and HG variant profiles of Phylotree. The narrowing down process involved the integration of the HGs with their MRCAs. QC analysis was included in this tool to check for possible artificial recombinations. We implemented the tool as described in HaploGrep 2 (a generic rule-based system). Once the variants and ranges of the fragment sequences had been obtained by the first tool, the second tool was used for HG tracking, rather than using the sequences. The third tool, “Conserved Region Map,” was developed based on the information of the variant positions of all the HGs presented by Phylotree (S5 Dataset).

### Accessory tools

Accessory tools were added to the server to help researchers obtain selected information about HGs, check for phantom mutants, convert variant profile to formats that are suitable for other web applications, and assign HGs using mtGenome sequences. The tool “HG Database” was constructed based on the tree structures of HGs and variant profiles of Phylotree. It helps researchers to explore an HG, its sub-HGs, and their differential variants. “Differentiation between HGs (HG Differentiation)” was designed to help search for differential variants among the user-inputted HGs. It was designed based on the Phylotree definitions. To analyze possible phantom mutations in a dataset (“Phantom Mutants”) and thus ensure the QC of the sample sequences, we implemented the tool following the methods used by HaploGrep 2. The tool “Haplogrouping with Complete mtGenome Sequences (MtGenome Haplogrouping)” was developed to definitively estimate the HG of a sample with complete mtGenome sequences. “Variant Format Conversion” was designed to enable researchers to conveniently convert variant profiles into different formats, including Phylotree, HaploGrep 2, EMPOP, and MitoTool. “Major HG-specific Variants” was included to show an outline of the phylogenic tree of major HGs and their specific variants defined by Phylotree.

### Comparison of web servers

We compared the haplogrouping performance of Haplotracker with that of MitoTool (Version Phylotree Build 17) and HaploGrep 2 (Version Phylotree Build 17) in the following tests using different mtDNA sequence sources. The haplogrouping data from MitoTool were obtained in June 2019, and the data from HaploGrep 2 and Haplotracker were obtained from June to September 2019. Significant differences between the HG estimation rates of the servers were determined using MedCalc Version 19.0.5. HG rankings of test samples were given by observing the positions of the HGs defined by Phylotree or determined using mtGenome sequences.

#### HG assignment test for complete Phylotree mtGenome sequences

In total, 8,216 complete mtGenome sequences from Phylotree were used, and the HG assignments from the servers were compared with Phylotree HGs.

#### Estimation test of Phylotree HGs with CR sequences only

CR sequences from all of the Phylotree mtGenome sequences above were used.

#### Further extensive evaluation

This included all available complete mtGenome sequences (n=48,086) from GenBank through to December 25, 2018. Of these, the sequences that were identically haplogrouped genome-wise by both HaploGrep 2 and Haplotracker were tested for subsequent HG estimation using their CR sequences (n=45,177). HaploGrep 2 and Haplotracker were compared.

#### Evaluation with new sequences

Additional GenBank mtGenome sequences (n=1,243) from December 26, 2018 to August 22, 2019 were tested. These sequences were not used for our haplotype DB construction and were obtained after the completion of Haplotracker development. Of these, 1,145 mtGenome sequences (genome-wise identically haplogrouped by both Haplotracker and HaploGrep 2) were selected and used for HG estimation according to their CR sequences.

#### Haplotracker HG estimation with HV region sequences

The HG estimation accuracy of Haplotracker using the HV region sequences (HV1, HV2, and HV3) was evaluated with the 8,216 mtGenome sequences from Phylotree.

### Laboratory-based application of Haplotracker

We evaluated the usefulness of Haplotracker’s haplogrouping in laboratory experiments with degraded DNA. We used an ancient DNA extract that had been reported previously (4). The sample DNA had been kept in a freezer for about three years after extraction. The sample (MNX3) was taken from 2000-year-old human skeletal remains from the Xiongnu Cemetery in Mongolia. The sample was from a West Eurasian male with mtDNA HG U2e1 and Y-SNP HG R1a1, and its CR sequences were directly used by Haplotracker. Laboratory experiments to obtain sequences of additional mtDNA fragments were conducted in the present study. Three CR sequence fragments (two HV1 and one HV2) were inputted into the main tool (Track by Sequence) of Haplotracker, and the HG was estimated.

To confirm the predicted HGs, we designed PCR primers for four fragments to identify coding-region variants that Haplotracker proposed for confirmation and differentiation (S8 Table). Conserved regions for PCR primers were searched using the additional Haplotracker tool (Conserved Region Map) by inputting the range of an approximately 400-bp-long region across the variant position and U2 for the HG target field. Multiplex primers for the coding-region variants were designed using LightCycler Probe Design Software 2.0 to avoid cross-complementarities between the multiple primers. Primer specificity in *Homo sapiens* was checked using primer-BLAST (https://www.ncbi.nlm.nih.gov/tools/primer-blast/). A LightCycler 96 (Roche) system was used for the real-time multiplex PCR amplification of the four coding-region fragments. The multiplex PCR reaction mixture (10 μl) consisted of 1X PCR buffer for Ex *Taq* HS (Takara), 0.3 mM dNTP (Takara), 0.3 μM of each primer (Macrogen), 2.5 mM MgCl_2_ (Roche), 0.6U Ex *Taq* HS, 0.5X SYBR Green I (Sigma), 1.5 mg/ml BSA (Life Technologies), PCR-grade water (Roche), and 3 μl of ancient DNA extract. The multiplex PCR cycling conditions were as follows: 1 cycle of 2 min at 95°C; 45 cycles of 10 s at 95°C, 30 s at 57°C, 40 s at 68°C, and 20 s at 72°C with a single fluorescence acquisition at the end of the extension step; and 1 melting curve cycle of 10 s at 95°C, 60 s at 65°C, and temperature increase from 65 to 95°C with a temperature transition rate of 0.1°C/s with continuous fluorescence acquisition at the rate of 5 readings/°C. The PCR products were diluted 500 times with PCR-grade water (Roche).

For the second round of real-time PCR (nested PCR), 2 μl of the diluted products was used for the template, and nested primers for each target coding-region fragment were added to the four separate reactions. The nested PCR reaction mixture (40 μl) consisted of 1X PCR buffer for Ex *Taq* HS (Takara), 0.2 mM dNTP (Takara), 0.2 μM of each primer (Macrogen), 2 mM MgCl_2_ (Roche), 0.25U Ex *Taq* HS, 0.5X SYBR Green I (Sigma), 0.5 mg/ml BSA (Life Technologies), PCR-grade water (Roche), and 2 μl of the diluted first-round product. The nested PCR cycling conditions were as follows: 1 cycle of 2 min at 95°C, 25 cycles of 15 s at 95°C, 15 s at 60°C, and 30 s at 72°C with a single fluorescence acquisition at the end of the extension step; and 1 melting curve cycle with the same conditions as described above. Amplicon identification and purity were estimated based on melting curve analysis. The PCR products were purified. Sequencing was conducted with the nested primers bidirectionally by Macrogen Incorporation in Korea. The sequences were read using SeqMan version 5.03 (DNASTAR) from the bidirectional sequence electropherograms of each DNA fragment. The sequences were applied to Haplotracker by adding them to the input fields next to the fragment sequences of HV1 and HV2 for confirmation of the predicted HG.

## Availability

Gotoh is an open-source C implementation of the Gotoh algorithm, a.k.a. Needleman-Wunsch with affine gap penalties (https://github.com/oboes/gotoh).

Phylotree Build 17 is the most recent version of the phylogenetic tree of global human mitochondrial DNA variation (https://www.phylotree.org).

MitoTool is a web-based bioinformatics platform providing a convenient, user-friendly interface for handling human mtDNA sequence data (http://www.mitotool.org/index.html).

HaploGrep 2 is a fast and free HG classification tool (https://haplogrep.i-med.ac.at).

EMPOP is a forensic mtDNA database (https://empop.online).

## Accession numbers

Because the number of mtDNA sequences used in this study is so large, their accession numbers are provided separately in S1, S3, and S7 Tables.

## Acknowledgements

We thank the following students at Chung-Ang University College of Medicine for their help in testing Haplotracker and the other web servers: Na-young Kim, Do-yoon Kim, Eo-jin Kim, Chae-won Kim, Sang-min Kim, Bum-seok Seo, Hae-won Shin, Kwang-wook Ahn, Yeol Yuh, Hee-jeong Lim, and Joong-hyun Cho. We also thank Jaehyeong Park at the Dept. of Computer Engineering, Seoul National University College of Engineering for his help in visual web design.

## Supporting information

**S1 Fig. Input interface for multiple sequence fragments of a sample**.

**S2 Fig. Input interface for multiple variant profiles of a sample**.

**S3 Fig. Example of detailed information of the HGs in Rank Group 1**.

**S4 Fig. Example of QC analysis of top-ranked HGs**.

**S5 Fig. Example of conserved region search for primer design**.

**S6 Fig. Comparison of HG predictions between servers**.

**S7 Fig. HG prediction results with the variant profile of a sample**.

**S1 Dataset. HGs and their variant profiles extracted from Phylotree mtDNA Build 17**.

**S2 Dataset. HG frequency carrying an extra variant in 118,869 haplotypes**.

**S3 Dataset. HG frequency carrying a missing variant in 118,869 haplotypes**.

**S4 Dataset. HG frequency in 118,869 haplotypes**.

**S5 Dataset. Variant positions and HGs at that position in Phylotree**.

**S1 Table. Comparison of servers with complete mtGenome sequences from Phylotree (n=8,216)**.

**S2 Table. Comparison of servers using only CR sequences from Phylotree (n=8,216)**.

**S3 Table. Comparison details for the servers using only CR sequences from Phylotree (n=8,216)**.

**S4 Table. Comparison of servers using CR sequences from GenBank (n=45,177)**.

**S5 Table. Comparison details for the servers using CR sequences from GenBank (n=45,177)**.

**S6 Table. Comparison of servers using CR sequences recently downloaded from GenBank**.

**S7 Table. Haplotracker’s HG prediction with HV region sequences from Phylotree (n=8,216)**.

**S8 Table. Primers and sequences used for the laboratory application of Haplotracker**.

